# A transcriptomic census reveals that Rbfox contributes to a broad but selective recapitulation of peripheral tissue splicing patterns in the thymus

**DOI:** 10.1101/2020.03.06.980870

**Authors:** Kathrin Jansen, Noriko Shikama-Dorn, Moustafa Attar, Stefano Maio, Maria Lopopolo, David Buck, Georg A. Holländer, Stephen N. Sansom

## Abstract

Thymic epithelial cells (TEC) control the selection of a T-cell repertoire reactive to pathogens but tolerant of self. While this process is known to involve the promiscuous expression of virtually the entire protein-coding gene repertoire the extent to which TEC recapitulate peripheral isoforms, and the mechanisms by which they do so, have remained largely unknown. We performed the first assembly-based transcriptomic census of transcript structures and splicing factor (SF) expression in mouse medullary TEC (mTEC) and 21 peripheral tissues. Mature mTEC expressed 60.1% of all protein-coding transcripts, more than was detected in any of the peripheral tissues. However, for genes with tissue-restricted expression, we found that mTEC produced fewer isoforms than did the relevant peripheral tissues. Analysis of exon inclusion revealed an absence of brain-specific micro-exons in mTEC. We did not find unusual numbers of novel transcripts in TEC and show that *Aire*, the facilitator of promiscuous gene expression, promotes usage of long transcripts but has only a limited impact on alternative splicing in mTEC. Comprehensive assessment of SF expression in mTEC identified a small set of non-promiscuously expressed SF genes amongst which we confirmed RBFOX to be present with AIRE in mTEC nuclei. Using a conditional loss of function approach, we show that *Rbfox2* promotes mTEC development and regulates the alternative splicing of promiscuously expressed genes. These data indicate that TEC recommission a small number of peripheral SFs, including members of the Rbfox family, to generate a broad but selective representation of the peripheral splice isoform repertoire.

## Introduction

T cells are essential for the generation and resolution of an adaptive immune response as they are uniquely able to distinguish between benign self and harmful non-self-antigens. Because T cell antigen receptor (TCR) specificity is generated pseudo-randomly, the functional utility and self-reactivity of TCRs must be screened during T cell development in the thymus (Klein et al. 2014). This process is critically dependent on different thymic stromal cells such as thymic epithelial cells (TEC) (Abramson and Anderson 2017). In an initial round of positive selection, cortical TEC (cTEC) positively select developing T cells (thymocytes) that express TCRs of sufficient affinity for self-MHC (Klein et al. 2014). Subsequently, both cTEC and mTEC deplete thymocytes that display TCRs with high affinity for self-antigens, a process designated thymocyte negative selection. In addition, mTEC divert a subset of self-reactive T cells to a natural T regulatory cell (nTreg) fate (Stritesky et al. 2012).

To achieve thymocyte negative selection, TEC express and present a molecular mirror of an individual’s self-antigens to maturing thymocytes. This process of “promiscuous gene expression” (PGE), is essential for the avoidance of autoimmunity and involves transcription of approximately 89% of protein-coding genes by the TEC population (Sansom et al. 2014; Abramson and Anderson 2017). The mechanisms by which TEC defy developmental and tissue-specific transcriptional controls to achieve PGE are only incompletely deciphered (Abramson and Goldfarb 2016), although the AutoImmune Regulator (Aire) (Sansom et al. 2014) and the transcription factor Fezf2 (Takaba et al. 2015) together with the chromatin remodeler Chd4 (Tomofuji et al. 2020) are known to enable PGE. Comprehensiveness of self-representation by TEC cannot, however, be measured by gene number alone because self-peptidome diversity is elaborated by alternative mRNA splicing (Matera and Wang 2014), expression of “untranslated” regions (Starck and Shastri 2016), RNA-editing (Danan-Gotthold et al. 2016), proteasome mediated splicing (Granados et al. 2015) and post-translational modifications (Raposo et al. 2018). In particular, alternative mRNA splicing greatly increases the complexity of the mammalian proteome: there are approximately three times more annotated protein-coding transcripts than genes in mice and humans (Zerbino et al. 2018).

The generation of alternative splice isoforms is tightly regulated during development and controlled by temporal and context-specific expression of SFs. The extent to which TEC reproduce peripheral splice isoforms is unclear but the transcriptome of TEC has been found to be unusually complex both in terms of isoform number (Keane et al. 2015) and splice-junction representation (Danan-Gotthold et al. 2016). These surveys, which compared TEC with limited numbers of peripheral tissues, reported that TEC reproduce a fifth of tissue-restricted isoforms (Keane et al. 2015) or only 20-60% of tissue-restricted splice junctions (Danan-Gotthold et al. 2016). Even less is known about the mechanisms in TEC that control alternative splicing. Because of its direct interactions with several splicing-related factors (Abramson et al. 2010), it was thought that AIRE might be involved in this process (Keane et al. 2015), but limited evidence supporting this idea suggests only a minor role (Danan-Gotthold et al. 2016). Instead, it is appealing to postulate that mTEC may reuse peripheral SFs to create tissue-specific splice variants. In one plausible model, mTEC might constitutively express a specific subset of peripheral SFs in order to achieve robust coverage of a limited repertoire of peripheral splice variants. In support of this possibility, an initial analysis identified seven RNA-binding factors with constitutive expression in murine mTEC (St-Pierre et al. 2015). In a second, alternative model, mTEC might employ promiscuously expressed SFs to achieve a broader stochastic coverage of peripheral isoforms. A possible side-effect of such a promiscuous splicing strategy would be the generation of spurious novel isoforms if nascent transcripts interact with SFs in TEC to which they are not exposed in the periphery.

Knowledge of splice isoform representation in the thymus is important for understanding central tolerance because it is known that the absence of tissue-specific splice isoforms can result in the development of pathogenic T cells able to incite autoimmunity (Klein et al. 2000). We therefore set out to perform the first comprehensive and unbiased census of isoform representation and SF expression in TEC and peripheral tissues. Our results show that TEC accurately reproduce a subset of tissue-specific splice isoforms in a process that involves the re-use of a small set of peripheral SFs. Finally, we demonstrate that members of the Rbfox SF family contribute to TEC development and help shape the generation of self-antigen splice-isoforms in these cells.

## Results

### TEC express fewer splice-isoforms per gene than peripheral tissues

To compare the splicing landscape of TEC with peripheral (i.e. non-thymic) tissues we performed deep stranded RNA-sequencing of immature and mature mTEC and integrated the data with that from similar sequencing of 21 separate tissues from the mouse ENCODE project (Pervouchine et al. 2015). To do so we constructed a mouse TEC and Tissue (mT&T) transcriptome assembly (Fig. 1A). This approach was essential because it was plausible that TEC might express large numbers of unknown transcript structures either through recapitulation of rare unannotated peripheral transcripts or as part of a stochastic program of alternative splicing. In agreement with previous reports (Keane et al. 2015; St-Pierre et al. 2015; Danan-Gotthold et al. 2016), we found that mature mTEC expressed a greater number of transcripts from protein-coding genes than any of the surveyed peripheral tissues (60.1%, Fig. 1B). However, modelling of the relationship between the number of detected genes and transcripts revealed that mature mTEC produced fewer transcripts per gene than was typically found in peripheral tissues (Fig. 1C).

**Figure 1:**
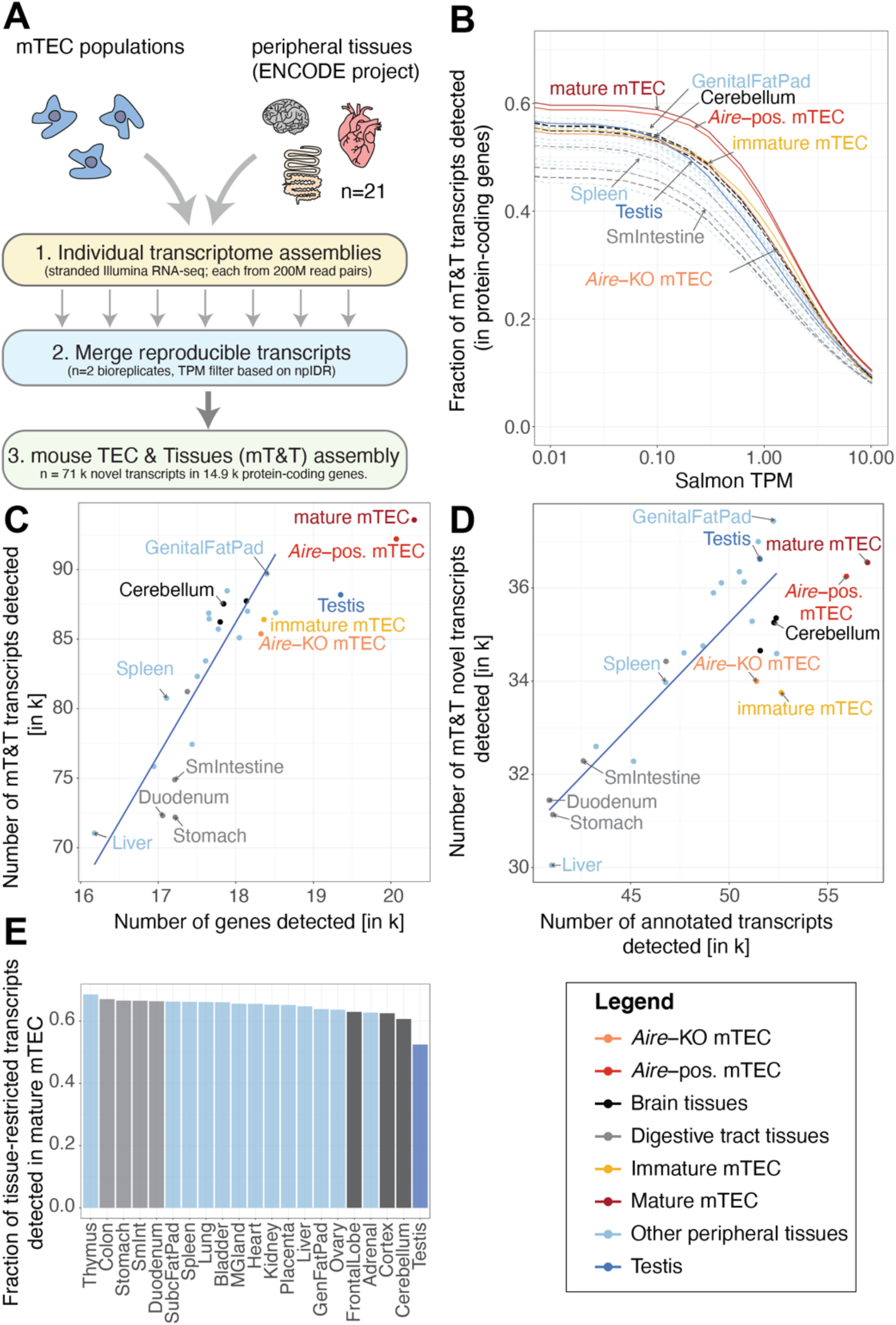
Comparative analysis of transcript expression in mTEC and peripheral tissues. (A) Generation of a common mouse mTEC and peripheral Tissues (mT&T) transcriptome assembly. (B) Fractions of transcripts from protein-coding genes detected in peripheral tissues and mTEC populations across a range of TPM thresholds. The scatter plots (C and D) show the relationships between the number of genes and transcripts (C) and the number of known vs novel transcripts (D) detected in peripheral tissues and mTEC populations. (E) Fractions of sets of tissue-restricted transcripts (*tau* ≥ 0.9) from peripheral tissues detected in mature mTEC. Analyses shown were restricted to protein-coding genes and performed using a single high-depth sample per tissue. Similar results were obtained using lower-depth biologically replicate sample pools (n=2, Supplemental Fig. 4). Trend lines in C and D were fitted to all samples except TEC and testis.

We next sought to determine the extent to which the splicing of promiscuously expressed genes in TEC recapitulates the patterns found in the normal tissue context. To do so we defined sets of *Aire*-regulated and non-*Aire* regulated “tissue-restricted antigen” (TRA) genes (Supplemental Methods and Supplemental Fig. 1). Relative to peripheral tissues we found that while TEC generated a similar fraction of isoforms from non-TRA genes they expressed a smaller fraction of isoforms from the promiscuously expressed TRA genes (Supplemental Fig. 2A-C). These observations did not appear to be a consequence of a lower sequencing coverage of promiscuously expressed genes in TEC (Supplemental Fig. 2D-F), because fewer isoforms were detectable in mTEC regardless of gene expression level (Supplemental Fig. 2G-I).

**Figure 2:**
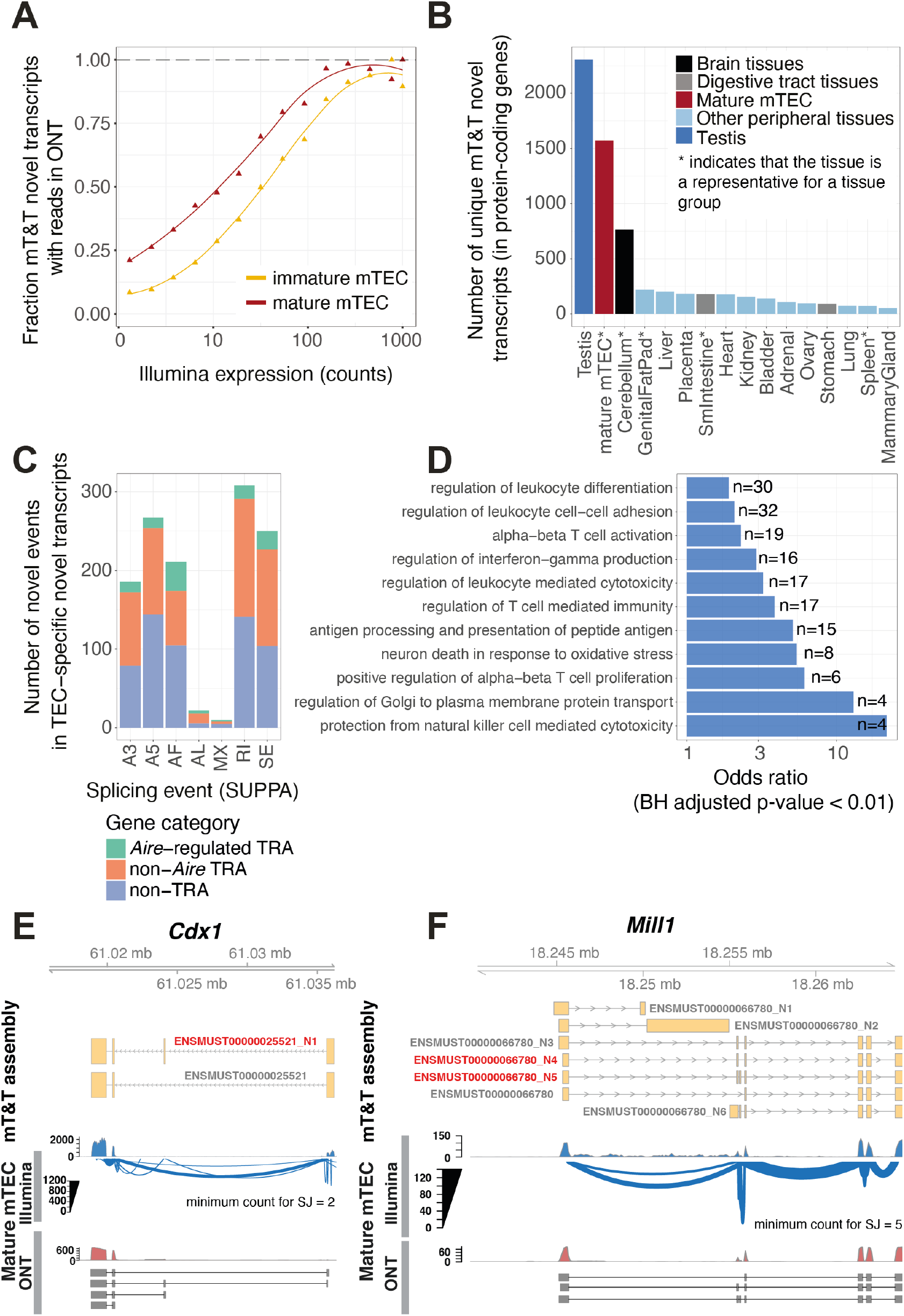
Identification and characterisation of TEC-specific novel transcripts. (A) Validation of novel mT&T transcripts using ONT RNA-sequencing. The fraction of novel transcripts supported by ONT reads are shown for mature mTEC (red) and immature mTEC (yellow). (B) Number of novel transcripts “uniquely” detected in mature mTEC and representative peripheral tissue samples (see Supplemental Methods and Supplemental Fig. 1A). (C) TEC-specific novel splicing events by event type and promiscuous expression status. Individual events may be counted in multiple categories. SE=skipped exon, RI=retained intron, MX=mutually exclusive exon, A3/A5=alternative 3’/5’ splice site, AF/AL=alternative first/last exon. (D) Selected GO biological processes significantly over-represented in the set of genes (n=1,167) from which the mTEC-specific novel transcripts were derived (one-sided FETs; BH adjusted p < 0.01). (E, F) *Cdx1* and *Mill1* are displayed as examples to demonstrate novel TEC-specific transcripts (red). Existence of the novel transcripts in mature mTEC was supported by both Illumina (sashimi plots) and long-read ONT (selected reads) data. Novel transcripts are indicated by the ‘_N’ suffix.

As previously reported (Keane et al. 2015; Danan-Gotthold et al. 2016), we found that the mature mTEC population co-expressed transcript isoforms which normally arise in distinct anatomical locations (Supplemental Fig. 3A-C). In addition, we found evidence that such transcripts can be produced by the same cell: single-cell RNA-sequencing data revealed that the thyroid and nervous-system specific transcript isoforms of *Calca* were often produced together in individual mTEC (Supplemental Fig. 3D). Finally, we assessed the representation of sets of tissue-restricted isoforms from peripheral tissues in mTEC. Transcripts with testis-restricted expression were most markedly under-represented in mTEC (52.5% detected), followed by those from the brain (60.7-62.9%), adrenal (62.7%) and ovary (63.6%) (Fig. 1E, with results from biological replicate sample pools shown in Supplemental Fig. 4).

**Figure 3:**
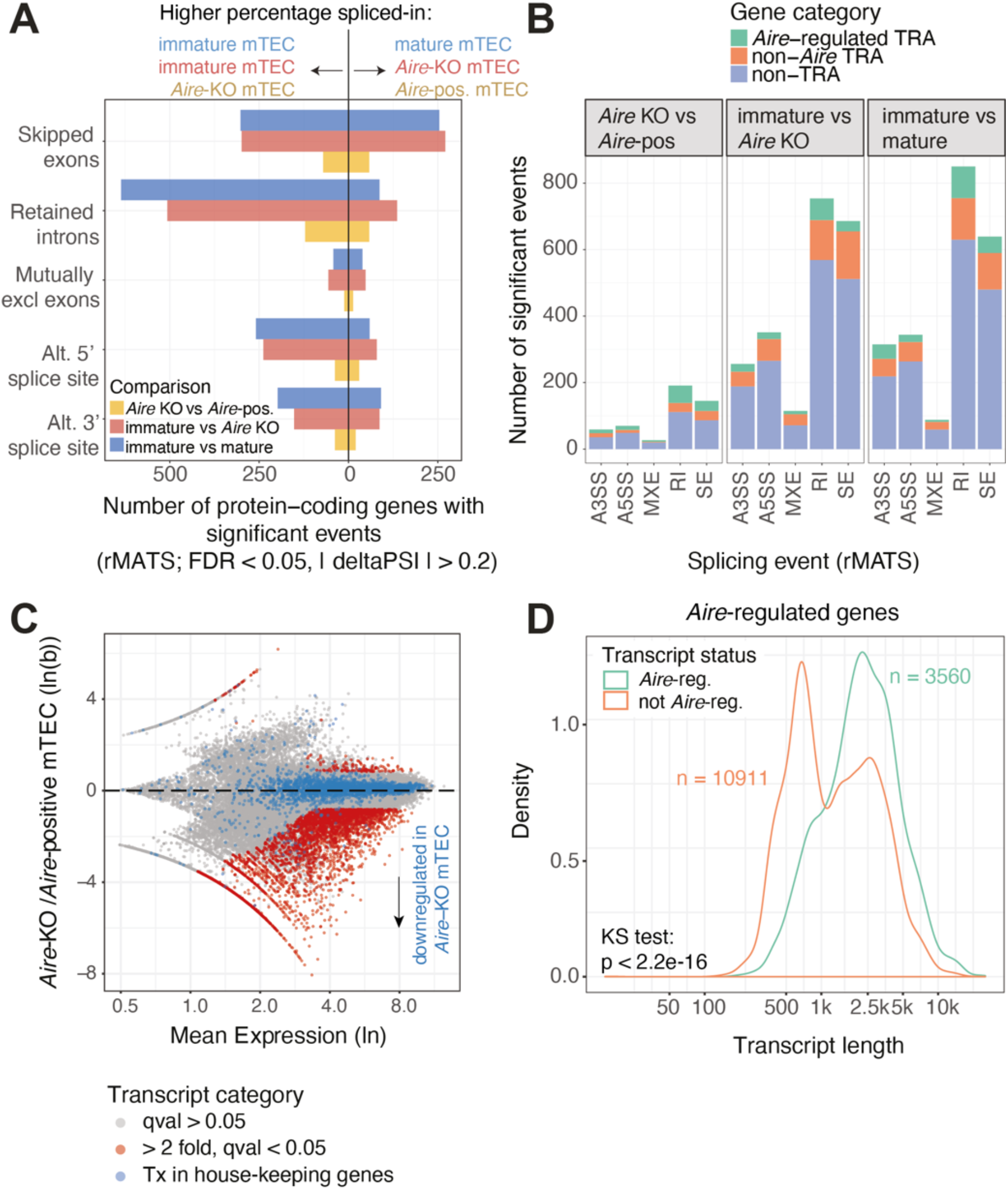
*Aire* promotes the generation of long transcripts in TEC. (A) The numbers of protein-coding genes (x axis) in which significant differential splicing events (y axis) were detected; comparisons of immature vs mature mTEC (blue), immature mTEC vs *Aire*-knockout mTEC (red) and *Aire*-knockout vs *Aire*-positive mature mTEC (yellow) (n=2 replicates per sample). (B) Breakdown of identified splicing events by event type and promiscuous expression status. SE=skipped exon, RI=retained intron, MXE=mutually exclusive exon, A3SS/A5SS=alternative 3’/5’ splice site. (C) MA plot of differential transcript expression in *Aire*-knockout compared to *Aire*-positive mTEC. Transcripts regulated by *Aire* are shown in red (Sleuth, Wald test, qval<0.05, fc ⪆2, n=2 replicates per sample). Transcripts from house-keeping genes are shown in blue. (D) The length distributions of *Aire*-regulated and non-*Aire*-regulated transcripts in *Aire*-regulated genes (analysis limited to genes that contained at least one significantly *Aire*-regulated transcript as defined in (C)).

**Figure 4:**
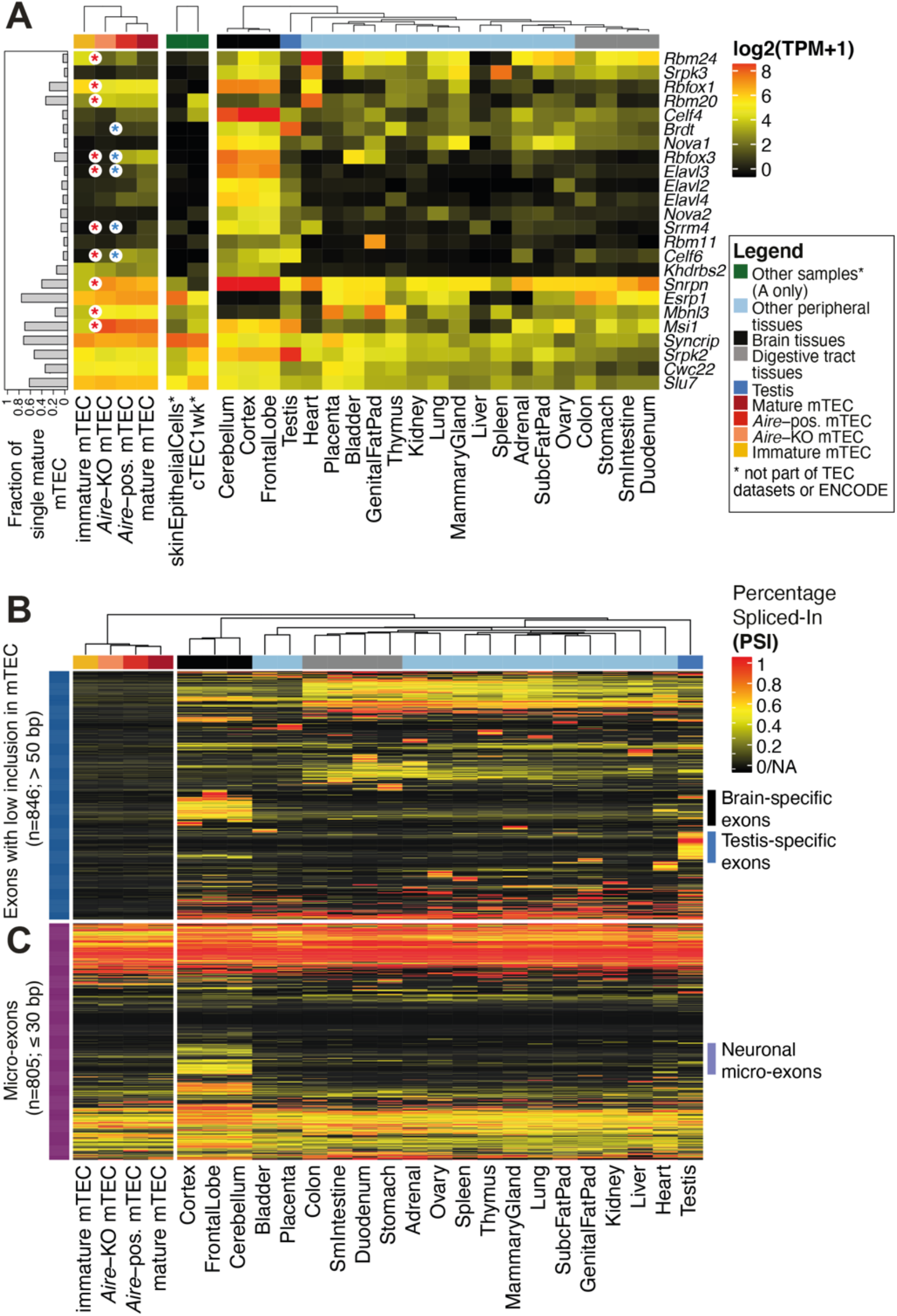
Selective expression of peripheral SFs and exons in mTEC. (A) Expression of a set of tissue-restricted (*tau* > 0.5) genes that encode for *bona fida* SFs (Supplemental Table 8) in mTEC populations, skin epithelia, cTEC and peripheral tissues (mouse ENCODE project). The bars (left) depict the fraction of a set of single mature mTEC that express each factor. Significant differences in expression between immature vs mature (*Aire*-KO) mTEC or *Aire*-KO vs *Aire*-positive mTEC are indicated by red and blue asterisks, respectively (BH adjusted p-value<0.05, | fc| >2, DESeq2 analysis of the population RNA-sequencing data, n=2 biological replicates/condition). (B) Patterns of protein-coding exon (>50bp in length) inclusion for exons with a low inclusion rate in mTEC (mean PSI<0.1; max PSI<0.2) that were included in at least one of the peripheral tissues (PSI>0.5) (C) Micro-exon (≤30bp) inclusion in transcripts from protein-coding genes in the mTEC population and peripheral tissue samples.

Finally, while mature mTEC expressed a higher number of known transcripts (n=57,019) than was observed in peripheral tissues, we found that they produced a relatively low number of novel transcripts (n=36,547, Fig. 1D). Together our results show that while, as a consequence of PGE, TEC produce an atypically high absolute number of transcripts, they tend to produce fewer isoforms per gene than is found in peripheral tissues.

### Genes harbouring novel transcripts in mTEC are associated with T-cell selection

Alternative splicing events are often associated with the evolution and modification of protein function (Kelemen et al. 2013). As the transcriptome of TEC is relatively understudied, we reasoned that novel splicing events specific to TEC might be associated with their specialised functions for T-cell selection. Using long-read Oxford Nanopore (ONT) sequencing (Supplemental Fig. 5) we validated the existence 64.3% of the mT&T novel transcript structures that were expressed at moderate or high levels in mature mTEC (>10 counts, Fig. 2A). Next, we identified novel transcripts that showed TEC or tissue-specific expression. For this analysis we chose a single tissue to represent each group of related tissues (Supplemental Fig. 1A). We found that mature mTEC expressed a higher number of unique novel transcripts than did brain tissue (as represented by the cerebellum), but substantially fewer than were found in the testis (Fig. 2B, Supplemental Table 3). The majority (85%) of the 1,572 novel transcripts uniquely expressed in mature mTEC were found for loci that do not require *Aire* for their expression (Fig. 2C).

**Figure 5:**
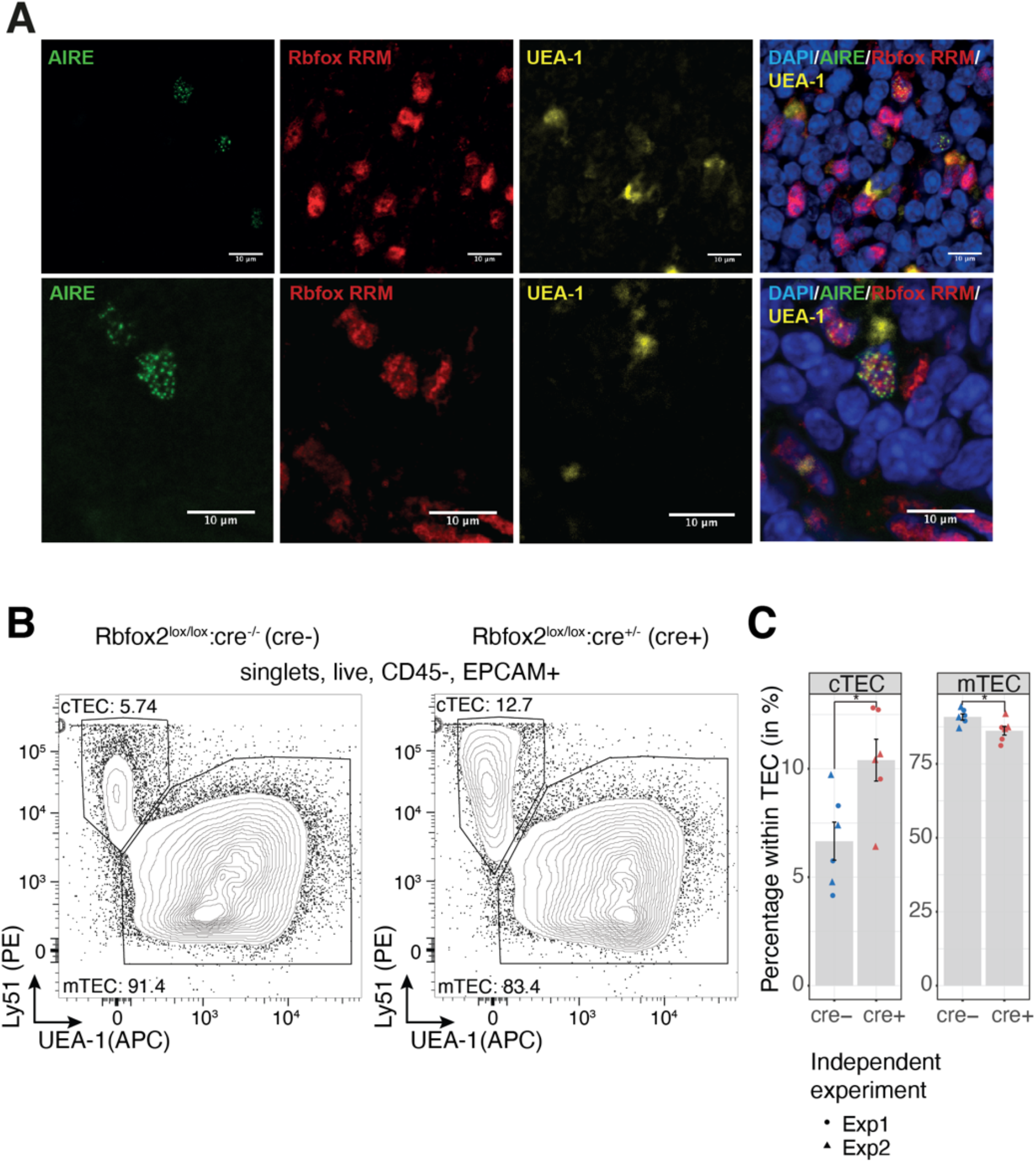
RBFOX is present with AIRE in mTEC nuclei and influences TEC development. (A) Confocal immunofluorescence analysis of the localisation of AIRE (green) and RBFOX (red) in the thymic medulla. RBFOX was detected using an anti-RRM domain specific antibody, mTEC were identified by reactivity with the lectin UEA-1 (yellow) and nuclei were labelled using DAPI (blue). Two representative sections are shown in the upper and lower panels. (B) Representative flow cytometric analysis of cTEC and mTEC frequencies amongst thymic epithelial cells extracted from *Rbfox2* tKO animals. (C) Quantification of cTEC and mTEC frequency in *Rbfox2* tKO animals. The mean ± SE from n=2 independent experiments is shown in bar graphs; * p-value<0.05 for two-sided Welch Two Sample t-test.

The 1,167 protein-coding genes which harboured TEC-specific novel transcripts displayed significant enrichments for Gene Ontology (GO) biological processes such as “regulation of leukocyte differentiation”, “regulation of T cell mediated immunity” and “antigen processing and presentation of peptide antigen” (Benjamini Hochberg (BH) adjusted p < 0.01, one-sided Fisher exact tests [FETs], Fig. 2D, and Supplemental Table 4) suggesting that they are likely to encode for non-promiscuously expressed factors that have a functional role in T-cell selection in TEC. Novel transcript structures were detected in genes of known importance for TEC function, including *Foxn1* (Zuklys et al. 2016) and *Aire* (Abramson and Goldfarb 2016) (Supplemental Table 3). Genes that produced transcripts harbouring novel exons or exon skipping events in TEC included *Cdx1*, a transcription factor linked with mTEC maturation (Handel et al. 2018) (Fig. 2E); *Cd80*, a receptor important for interaction with T-cells via CD28 and CTLA4; the MHC class II gene H2-Aa; *Mill1*, an MHC class I-like molecule that is known to be expressed on a subpopulation of TEC (Kajikawa et al. 2006) (Fig. 2F) as well as Skint family members, including *Skint2* which has been reported to be a novel negative T cell regulator (Yang et al. 2007).

Together these observations suggest that the novel transcript structures detected in thymic epithelial cells may have relevance for TEC function.

### *Aire* contributes to alternative splicing and promotes expression of long transcripts

To clarify the role of Aire in alternative splicing in TEC, we also generated deep stranded Illumina sequencing data from mature *Aire*-positive and mature *Aire*-knockout mTEC. We discovered 492 significant (5% FDR, rMATS) *Aire*-regulated alternative splicing events in transcripts from 459 protein-coding genes (Fig. 3A, Supplemental Table 5). There was, however, a much larger difference in transcript splicing between immature and mature wildtype mTEC (n=2,236 events in n=1,967 protein-coding genes, 5% FDR, rMATS analysis, Fig. 3A and Supplemental Table 5). Analysis of the differences in splicing events between immature wildtype and mature *Aire*-knockout mTEC confirmed a relatively limited contribution of *Aire* to alternative splicing (Fig. 3A). Of note, the majority of the *Aire*-controlled and TEC maturity-related alternative splicing events were found in transcripts encoding non-TRA genes (Fig. 3B). In addition, a large number retained introns were found in the transcripts of immature mTEC (n=760, Fig. 3A). Coordinated changes in intron retention during cellular differentiation are not unusual, and transcripts harbouring retained introns can arise from genes that have specialised cellular functions (Jacob and Smith 2017).

AIRE is known to promote expression of distal exons (Meredith et al. 2015) and the release of stalled polymerases (Giraud et al. 2012). We therefore investigated whether *Aire* might favour the production of long transcripts. Differential transcript usage analysis (Fig. 3C) revealed that *Aire* positively regulated the generation of a large number of transcripts (n=4,027, BH adjusted p < 0.05, fold change ⪆2; Supplemental Table 6). We found that *Aire*-regulated transcripts arising from *Aire*-regulated genes had a significantly different length distribution (p=2.2 ×10^−16^ Kolmogorov-Smirnov test) being on average over 1kb longer than their non-*Aire*-regulated counterparts (Fig. 3D). These findings show that *Aire* plays a major role in promoting the production of long transcripts while only having a limited impact on the generation of alternative splicing events. Hence, splicing factors other than *Aire* must be primarily responsible for the alternative splicing of transcripts in mature mTEC.

### Medullary TEC express a distinct set of SFs that includes *Rbfox1*

The ability of mTEC to express a large number of peripheral isoforms suggested that they might re-use tissue-specific SFs. To investigate this possibility, we systematically identified a set of 146 splicing related genes that showed tissue-restricted expression (Supplemental Methods, Supplemental Figure 6 and Supplemental Table 7). Based on literature searches we further narrowed our focus to n=24 tissue-restricted splicing factors (TRSF) with known roles in controlling alternative splicing (Fig. 4A, Supplemental Table 8). Of these we noted the frequent (detected in >20% of single mTEC) and non-promiscuous expression of *Rbfox1, Rbm20* and *Msi1* in mature mTEC. These factors also showed little, if any, expression in skin epithelia (Fig. 4A) suggesting that they may be relevant to the specialised function of mTEC. Rbfox1 and Rbm20 control alternative splicing in the brain and myocardium (Guo et al. 2012; Conboy 2017) while Msi1 is an RNA binding protein implicated in regulating splicing in photoreceptors (Murphy et al. 2016). Meanwhile, we noted that the majority of the 24 TSRFs −including those restricted in their expression to the brain (such as *Nova1, Nova2, Elav2, Elav3, Elav4* and *Rbfox3*), testis (*Brdt*) and heart (*Rbm24*) - were expressed at low frequency or showed only promiscuous expression in mature mTEC (Fig. 4A).

**Figure 6:**
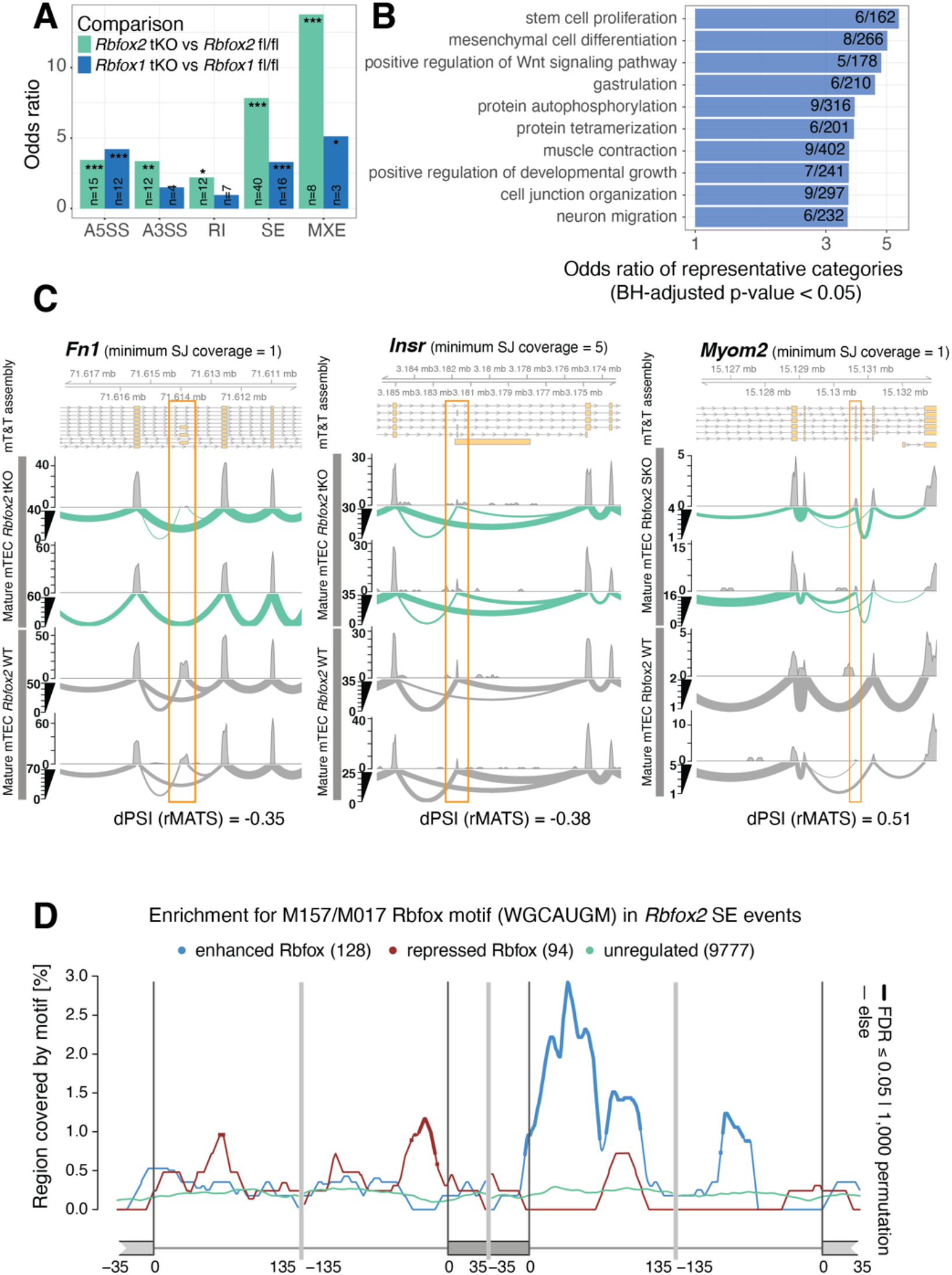
Rbfox regulates alternative splicing in TEC. (A) The barplots show the enrichments (ORs) of previously predicted Rbfox target genes (Weyn-Vanhentenryck et al. 2014) in the sets of genes differentially spliced in the *Rbfox1* tKO or *Rbfox2* tKO mature mTEC (Two-sided FETs, BH adjusted p-values). * p<0.05, ** p<0.01, *** p<0.001; abbreviations as for Fig 3B. (B) Selected GO biological processes significantly over-represented (one-sided FETs, BH adjusted p-values < 0.05) in genes that contained significantly Rbfox2 regulated SE events in mature mTEC. (C) Three examples of significantly (FDR<0.05) Rbfox2 regulated splicing events in mature mTEC. *Fn1* and *Insr* are known Rbfox target genes (Chen and Manley 2009; Weyn-Vanhentenryck et al. 2014). *Myom2* is an example of a non-*Aire* TRA gene. (D) Enrichment of the Rbfox recognition motif (M159/M017)(Ray et al. 2013) in the sequences surrounding exons that were found to be significantly regulated by Rbfox2 in mature mTEC (Supplemental Fig. 12B). The lines show enrichments for sets of exons found to be enhanced (blue), repressed (red) or not significantly regulated (green) by Rbfox2. Thicker lines indicate regions of statistically significant enrichment (FDR≤0.05, n=1,000 permutations).

In line with the absence of neural and testis-specific SFs, sets of coding exons frequently included in transcripts detected in the brain and testis were found to be excluded from mTEC mRNA (PSI < 0.1, Fig. 4B). In particular, we noted that brain specific micro-exons (≤ 30 bp) were rarely included in mTEC transcripts (Fig. 4C). This observation may be explained by the low expression of *Srrm4* in mTEC (Fig. 4A) as this factor is known to be responsible for promoting the inclusion of neuronal micro-exons (Quesnel-Vallieres et al. 2015; Quesnel-Vallieres et al. 2016). In summary we found that mTEC constitutively express only a small number of peripheral SFs. Together with the clear absence of a set of neuronal micro-exons, these results suggest that mTEC may have evolved to selectively represent specific facets of the peripheral tissue isoform repertoire.

### RBFOX is present with AIRE in mTEC nuclei and influences TEC development

Amongst the TRSFs expressed by mTEC, *Rbfox1* was of particular interest due to the established roles of Rbfox in controlling alternative splicing during muscle and brain development (Conboy 2017). We noted that *Rbfox1* and *Rbfox2* transcripts in mTEC predominantly included the neuronal B40 exon and excluded the muscle-specific M43 exon (Damianov and Black 2010) (Supplemental Fig. 7). Confocal image analysis of AIRE-expressing mTEC with a pan Rbfox anti-RRM domain antibody revealed that Rbfox factors are present in the nucleus in proximity to AIRE speckles (Fig. 5A).

Both *Rbfox1* and *Rbfox2* showed a robust non-promiscuous and tissue-restricted expression in mTEC (Fig. 4A, Supplemental Fig. 7). To investigate their functions in mTEC, we conditionally knocked them out (Gehman et al. 2011; Gehman et al. 2012) using the expression of Cre recombinase under the transcriptional control of the *Foxn1* locus (Zuklys et al. 2009). Total thymus cellularity of *Rbfox1* thymus knockout (tKO) (*Rbfox1*^*lox*/lox^:*Foxn1*^*cre*/+^) mice was 15% increased whereas that of age-matched *Rbfox2* tKO (*Rbfox2*^*lox/lox*^*:Foxn1*^*cre/+*^) mice remained unchanged when compared to controls (Supplemental Figs 8C, 9C). However, the phenotype of cTEC, mTEC, immature mTEC, and mature mTEC was unaffected in *Rbfox1* tKO mice (marker phenotypes in Supplemental Table 9, Supplemental Fig. 8). In contrast, *Rbfox2* tKO mice showed an overall reduction in TEC frequency (CD45-EpCAM+, *Rbfox2* tKO 0.12 % vs WT 0.17 % of all thymic cells recovered, p<0.05, two-sided Welch Two Sample t-test, Supplemental Fig. 9D-E). The *Rbfox2* tKO mice also had proportionally more cTEC (10.4% vs 6.7% of all TEC, p=0.02, Welch Two Sample t-test) and proportionally fewer mTEC (86.2% vs 90.8% of all TEC, p=0.03, Welch Two Sample t-test) present in their epithelial scaffolds (Fig. 5B-C). *Rbfox1/2* tKO (*Rbfox1*^*lox/lox*^: *Rbfox2*^*lox/lox*^:*Foxn1*^*cre/+*^) animals showed a TEC population composition that was quantitatively similar to that of *Rbfox2* tKO mice (Supplemental Fig. 10) suggesting that, as in other tissues (Gehman et al. 2011), *Rbfox2* can compensate for the loss of *Rbfox1* in TEC. Thymocyte development and positive selection were quantitatively normal in the Rbfox1/2 tKO animals (Supplemental Fig. 11). In summary these data demonstrate that *Rbfox1* contributes to the regulation of thymic cellularity and that *Rbfox2* increases the relative frequency of mTEC and decreases the relative frequency of cTEC within the TEC compartment.

### Rbfox contributes to the alternative splicing of self-antigen transcripts in mTEC

To investigate the impact of Rbfox on the mTEC transcriptome we performed RNA-sequencing of immature and mature mTEC isolated from *Rbfox1* tKO, *Rbfox2* tKO and control mice. In mature mTEC loss of *Rbfox1* or *Rbfox2* induced only minor changes in gene expression (Supplemental Fig. 12A) but caused hundreds of alterations in alternative splicing (*Rbfox1* tKO: n=559 events in 535 genes; *Rbfox2* tKO: n=668 events in 624 genes; FDR <0.05, | delta PSI| >0.02, Supplemental Fig. 12, Supplemental Tables 10, 11). In mature mTEC, *Rbfox1* and *Rbfox2* controlled the alternative splicing of 123 events in 104 TRA genes and 122 events in 110 TRA genes, respectively. In immature mTEC *Rbfox1* and *Rbfox2* exerted similar effects but controlled the alternative splicing of a smaller number of TRA genes (72 events in 51 TRA genes and 43 events in 36 TRA genes, respectively) in keeping with the lower level of promiscuous gene expression in these cells (Supplemental Fig. 13 A-C). The splicing of 10 genes was regulated by both *Rbfox1* and *Rbfox2* in mature mTEC (odds ratio [OR]=4.64, p=1.71 ×10^−4^, two-sided FET) while the splicing of 6 genes was regulated by both factors in immature mTEC (OR=18.5, p=3.23×10^−6^, Supplemental Fig. 13).

We next sought to establish whether the genes spliced by Rbfox in mature mTEC included those previously predicted to be Rbfox targets in the mouse brain (Weyn-Vanhentenryck et al. 2014). In mature mTEC we identified large and significant overlaps between predicted Rbfox target genes and those for which Rbfox2 controlled (i) exon skipping in mTEC (OR=7.8, BH adjusted p=5.4×10^−19^) or (ii) the use of mutually exclusive exons (OR=13.8, BH adjusted p=1.4×10^−5^) in mTEC (Fig. 6A). In keeping with this observation, many of the TRA genes alternatively spliced by Rbfox2 showed specific expression in neuronal tissues (Supplemental Fig. 12D). GO biological processes over-represented amongst the genes alternatively spliced by Rbfox2 in mature mTEC included “muscle contraction” and “neuron migration” in line with the known roles of Rbfox family members in muscle and neuronal tissues (Conboy 2017)(Fig. 6B). Examples of genes harbouring Rbfox2 mediated alternative splicing events in mature mTEC included *Fn1* and *Insr*, two known Rbfox2 target genes (Chen and Manley 2009; Weyn-Vanhentenryck et al. 2014) as well as the TRA *Myom2* (Fig. 6C).

Previous studies have identified an enriched Rbfox recognition motif in proximity to Rbfox regulated exons (Weyn-Vanhentenryck et al. 2014). We observed a significant enrichment of the conserved Rbfox RRM WGCAUGM motif (Ray et al. 2013) upstream of exons repressed and downstream of exons enhanced by *Rbfox2* in both mature and immature mTEC (Fig. 6D, Supplemental Fig. 13D), following the patterns previously described for this factor (Weyn-Vanhentenryck et al. 2014). In summary these data provide evidence that Rbfox SFs directly regulate the splicing of both promiscuously and non-promiscuously expressed genes in mature mTEC.

## Discussion

Our work shows that while mature mTEC produce an exceptionally high proportion (approximately sixty percent) of peripheral splice variants they are unable to recreate the full diversity of isoforms that are present in the periphery. The concept of selective representation of peripheral isoforms in TEC is supported by a recent qPCR-based study which estimated that a quarter of the genes studied contained epitopes hidden from the thymus (Shilov et al. 2019). Overall, our rigorous, transcriptome-assembly based census of transcript structures showed a remarkably even representation of tissue-restricted transcripts from the 21 surveyed peripheral tissues in mTEC. Most notably we found a lower representation of transcripts from the testis and brain together with a corresponding absence (or only very weak) expression of brain and testis-specific SFs in mTEC. It is possible that it may be functionally less important to educate T cells against splice-isoform epitopes specific to these tissues, which constitute so-called immune privileged sites. Further, the unusually high complexity of splicing in the testes and brain suggests the possibility that immune surveillance may act as a constraint on the evolution of splicing complexity in non-immune privileged peripheral tissues. Our finding that neuronal micro-exons are not frequently spliced into transcripts in mTEC is likely explained by the absence of the micro-exon SF *Srrm4* in these cells (Quesnel-Vallieres et al. 2015). The limited inclusion of neuronal micro-exons in mTEC provides a possible link between the observations that such exons are mis-regulated in the brains of patients with autism (Irimia et al. 2014), and growing evidence that autoimmunity may be involved in autism (Hughes et al. 2018). Aside from the testis and cerebellum, adrenal and liver tissues showed a weaker isoform representation in mature TEC. In humans, these tissues are known sites of autoimmunity: autoimmune adrenalitis is the most common cause of Addison’s disease, and liver diseases such as autoimmune hepatitis, primary biliary cirrhosis and sclerosing cholangitis are all thought to be the consequence of autoimmunity (Decock et al. 2009). Alternative splicing has been implicated in such diseases (Webster 2017), and our data suggests that lack of thymic representation of isoforms specific to these tissues might contribute to a higher susceptibility for autoimmunity.

Our investigations revealed a large number of splicing differences between immature and mature mTEC. One possible explanation for this observation is the expression of *Aire* in mature mTEC but our analysis of *Aire*-knockout mTEC confirms that this molecule plays only a small role in alternative splicing in mTEC. Rather, we found that *Aire* does promote the use of long transcripts, which is in line with the concept that it releases stalled RNA polymerases (Giraud et al. 2012). A comprehensive survey of splicing factor expression revealed that mTEC complement typical epithelial splicing factors such as *Espr1/2* with a small number of peripheral restricted splicing factors. Unlike skin epithelia, mature mTEC showed non-promiscuous expression of *Rbm20, Msi1* and *Rbfox1*. This observation, along with the apparently incomplete representation of peripheral structures and absence of excessive numbers of novel splice-isoforms in mTEC, suggests that TEC undertake a selective program of alternative splicing to ensure the accurate representation of a subset of the peripheral splice isoform repertoire. We hence conclude that transcript structures in mTEC are primarily shaped by a small number of splicing and mRNA processing factors (Guyon et al. 2020) and that mTEC do not rely on “promiscuous” mechanisms to promote and increase splice isoform diversity, as previously suggested (Keane et al. 2015). Rather (and assuming that our observations hold in humans), a limited thymic representation of peripheral splice isoforms is consistent with the concept that tissue-specific isoforms are relevant sources of auto-antigens in immune-mediated diseases (Evsyukova et al. 2010; Juan-Mateu et al. 2016; Newman et al. 2017). Furthermore, characterisation of transcript structures in human thymic epithelial cells would be expected to aid the identification of autoantigen transcripts encoding epitopes against which central tolerance has not been achieved (Ng et al. 2004). Knowledge of such epitopes is of great value for development of antigen-specific therapies for autoimmune disease such as those based on the use of tolerizing peptides or tolerogenic dendritic cells (Pozsgay et al. 2017; Mosanya and Isaacs 2019).

Our discovery of non-promiscuous *Rbfox1* expression in TEC was of particular interest because this factor is otherwise restricted to muscle and neural tissues in which it plays important developmental roles (Conboy 2017). We detected *Rbfox1* in medullary but not cortical TEC suggesting that it may be important for the development or function of this subpopulation. We therefore performed functional analysis of the role of *Rbfox1* and its homologue *Rbfox2* in mTEC, excluding *Rbfox3* from our investigations as it showed only weak and promiscuous expression in these cells. Phenotypic analysis of TEC in animals with a selective loss of one or both of these factors showed that *Rbfox2* increases mTEC frequency and decreases cTEC frequency within the TEC population. Recently, it has been shown that whilst in the embryonic and new-born thymus cortical and medullary TEC arise from a common bipotent progenitors, in the adult mouse mTEC are mostly replenished by lineage-restricted cells (Ohigashi et al. 2015). While our data suggest that *Rbfox* might influence the cell fate choices of the bipotent progenitors or act later to promote the differentiation within the mTEC lineage other possibilities cannot be excluded.

In mature mTEC we found that Rbfox factors shape the splicing of both promiscuously and non-promiscuously expressed genes. The larger role identified for *Rbfox2* in this process was not unexpected, as, in the cerebellum it is known that while *Rbfox2* can largely compensate for loss of *Rbfox1, Rbfox1* is less well able to ameliorate an absence of *Rbfox2* (Gehman et al. 2012). In addition, the changes in alternative splicing identified following loss of *Rbfox2* may involve *Rbfox1* as *Rbfox1* was itself differentially spliced in the absence of *Rbfox2*. We found examples of both *Aire*-dependent and *Aire*-independent TRA that were alternatively spliced by *Rbfox2* in mTEC. Given the weak expression of TRA in mTEC and the relatively small amount of biological material sequenced for this analysis, we expect the actual number of TRA regulated by Rbfox factors in mTEC to be substantially higher than that reported here. In summary, our data suggest that mTEC re-use a small set of peripheral SFs that includes Rbfox in order to selectively reproduce a broad subset of the peripheral splice isoform repertoire.

## Methods

### Mice

Wildtype C57BL/6 mice were obtained from Harlan Laboratories or Janvier and maintained as a laboratory in-house colony. *Aire*^*GFP/+*^ mice were previously described (Sansom et al. 2014). *Rbfox1* and *Rbfox2* mutant mice (Gehman et al. 2011; Gehman et al. 2012) were maintained on a mixed 129S2/Sv x C57BL/6J background.

### Isolation, sorting and immunostaining of thymic epithelial cells

TEC were isolated from multiple thymi and sorted by surface phenotype (Supplemental Table 9 and Supplemental Methods) (Dhalla et al. 2020). Sections were prepared and stained for confocal microscopy as described in the Supplemental Methods.

### RNA-sequencing data

For the analysis of immature, mature and *Aire*-knockout mTEC poly(A)+ RNA-seq libraries were prepared from cells pooled from multiple mice (1 μg of total RNA; 4 weeks old; n=2 biological replicates, Supplemental Table 1) and subjected to 101 bp paired-end stranded Illumina RNA-seq. For the analysis of *Rbfox1* tKO and *Rbfox2* tKO animals, immature and mature mTEC were isolated from individual mice and littermate controls (10-15k cells per animal, n=2 biological replicates, 4 weeks old). Stranded RNA-seq libraries were prepared (NEB) and subjected to 150bp paired-end Illumina HiSeq 4000 sequencing. For long-read sequencing CD80+86+ or CD80-86-mTEC were obtained from 4-6 week old female wildtype mice and processed as above. Details of the ONT sequencing can be found in the Supplemental Methods.

### Computational methods

The computational methods are described in the Supplemental Methods.

### Data access

RNA-sequencing data are available via NCBI Gene Expression Omnibus accession GSE145931.

## Acknowledgements

We thank Prof. Chris Ponting (University of Edinburgh) for support and advice. The anti-RRM antibody and Rbfox mutant animals were kind gifts of Prof. Douglas Black and Julia Nikolic (University of California, Los Angeles). We thank Prof. Black for helpful comments and advice. This work was initiated with funds from the Medical Research Council (MRC) CGAT programme [G1000902]. SNS and GAH were supported by funding from the Wellcome Trust (#066521). SNS and MA are supported by funding from the Kennedy Trust for Rheumatology Research (KTRR). KJ was supported by a Wellcome Trust PhD studentship and funds from the MRC (MR/S025308/1, MR/S035850/1).

## Author contributions

K.J. performed the experiments and computational analyses. N.S.D. generated RNA-sequencing data from the wildtype and *Aire*-knockout mTEC. S.M. assisted with the flow cytometry experiments. M.A. and M.L. performed, and D.B. was responsible for, the ONT library construction and sequencing. S.N.S. conceived the study with input from G.A.H. K.J., G.A.H. and S.N.S. designed the experiments and analyses, interpreted results and wrote the manuscript. G.A.H. and S.N.S. supervised the study.

